# Luciferase Immunoprecipitation Systems immunoassay is a sensitive, rapid method to detect allergen component-specific IgE

**DOI:** 10.1101/2020.01.24.917153

**Authors:** Adora A. Lin, Natalia S. Perez, Pamela A. Frischmeyer-Guerrerio, Thomas B. Nutman

## Abstract

Current assays to detect allergen-specific IgE have constraints related to obtaining pure, conformationally active allergen, variability in allergen extracts, sample volume required, and turnaround time. The luciferase immunoprecipitation systems (LIPS) immunoassay is a fast, sensitive assay created using recombinant antigens that requires a low specimen volume. These assays can also be easily modified to detect multiple antigens and antibody isotypes. Here, we demonstrate the use of LIPS assays as an innovative method to quantitatively measure allergen component-specific IgE in small sample volumes. Sera from healthy volunteers, helminth-infected adults, and peanut-allergic children were screened for IgE to cat using ImmunoCAP. These samples were also measured for IgE against Fel d 1 using LIPS. LIPS signal correlated to cat IgE levels with *r*_S_ = 0.6204, p < 0.001. The LIPS signal: noise ratio differed significantly between cat IgE-samples and cat IgE+ samples with values > 0.5 kU/L, with the ability to differentiate cat IgE – individuals from cat IgE+ individuals with 85% sensitivity and 76% specificity. Given their rapidity, efficiency, sensitivity, and quantitation over a broad dynamic range, LIPS immunoassays can be a robust and flexible tool with potential uses in allergy research, diagnostics, and treatment.

## Introduction

Current laboratory methods to determine the presence of circulating allergen-specific IgE involve the use of crude biological extracts or purified recombinant allergens.^1^ Extracts may differ in composition due to difficulties in obtaining a pure, raw material, and/or differences in processing,^1, 2^ and recombinant allergens are difficult to obtain in biologically relevant, conformationally active forms.^2.^ These limitations with the allergen can lead to variable results in assays.^1^ In addition, challenges arise in patients undergoing evaluation for multiple allergens with restrictions on blood draw volume, such as children. All of these constraints indicate a need for an efficient and sensitive assay using small sample volumes and recombinant proteins. The luciferase immunoprecipitation systems (LIPS) immunoassay has been shown to be an effective method of detection of autoantibodies and antibodies against infectious agents with high sensitivity and specificity.^3^ Advantages of this method over other assays for antibody detection include fast turnaround time, the use of recombinant antigens without elaborate purification schemes, and ease of optimization.^3^ Here, we demonstrate the power of synthetic biology coupled with the LIPS immunoassay as a novel, rapid method to detect allergen-specific IgE in low-volume samples.

## Methods

### Clinical samples

Sera were obtained from healthy volunteers (n=76), helminth-infected adults (n=26), and peanut-allergic children (n=12). All sera were obtained under NIH IRB-approved protocols (99-CC-068, 88-I-83 and15-I-0162), and written informed consent was obtained from all subjects or their parents/legal guardians. Samples were screened for IgE to cat using ImmunoCAP. Cat IgE levels ranged from 0 to >100 kU/L.

### LIPS assay

LIPS immunoassays were performed as described previously.^3^ Briefly, 5 µl of patient sera were combined with an optimized amount (∼1 million light units [LU]) of *Renilla* luciferase-Fel d 1 fusion protein extracts in a 96-well microtiter plate and incubated for 10 minutes. Resulting IgE-fusion protein complexes were immunoprecipitated by incubation with anti-IgE beads in a 96-well filter plate. Following several washes to remove unbound fusion proteins, LU were measured on a Berthold LB 960 Centro luminometer (Berthold Technologies, Oak Ridge, TN) using a coelenterazine substrate mixture (Promega, Madison, WI). All samples were run in duplicate.

### Fusion protein constructs

*Renilla* luciferease fusion protein constructs were synthesized for the Fel d 1 component of cat allergen by GenScript Biotech (Piscataway, NJ) using the amino acid sequence previously published ^10^. Codons were optimized for expression in mammalian cells. Constructs were transfected into FreeStyle™ 293-F cells (ThermoFisher, Waltham, MA). Fusion proteins were isolated after 48-72 hours of cell culture by cell lysis in lysis buffer composed of 50 mM Tris, pH 7.5, 100 mM NaCl, 5 mM MgCl2, 1% Triton X-100, 50% glycerol and protease inhibitors. Anti-IgE beads were created for use in the LIPS assay by coupling goat anti-human IgE (SeraCare KPL, Gaithersburg, MD) to Ultralink beads (ThermoFisher, Waltham, MA).

### Statistical analyses

Figures and statistical analyses, Receiver Operating Characteristic (ROC) curve analysis, and correlations (Spearman’s rank), were done using Prism 6.0 (GraphPad Software, Inc., San Diego, CA). The nonparametric Mann-Whitney test was used to estimate differences in LIPS signal between two groups.

## Results

For Fel d 1 LIPS, there was no significant difference between the signal: noise ratio for cat IgE – samples and cat IgE+ samples with values of 0.36-0.49 kU/mL. However, the signal: noise ratio differed significantly between cat IgE-samples and cat IgE+ samples with values > 0.5 kU/mL and encompassed a broad dynamic range (Figure 1A, *p* < 0.02). LIPS signal correlated to cat IgE levels with *r*_S_ = 0.6204, *p* < 0.001(Figure 1B). ROC analysis gave an area under the curve of 0.7929 (*p* < 0.001), which increased to 0.8455 (*p* < 0.001) if cat IgE+ samples were defined as being ≥ 0.5 kU/mL. A threshold of 1647 LU/µl distinguished cat IgE –individuals from cat IgE+ individuals with 85% sensitivity and 76% specificity. Assays were completed in less than one hour.

**FIGURE 1.**
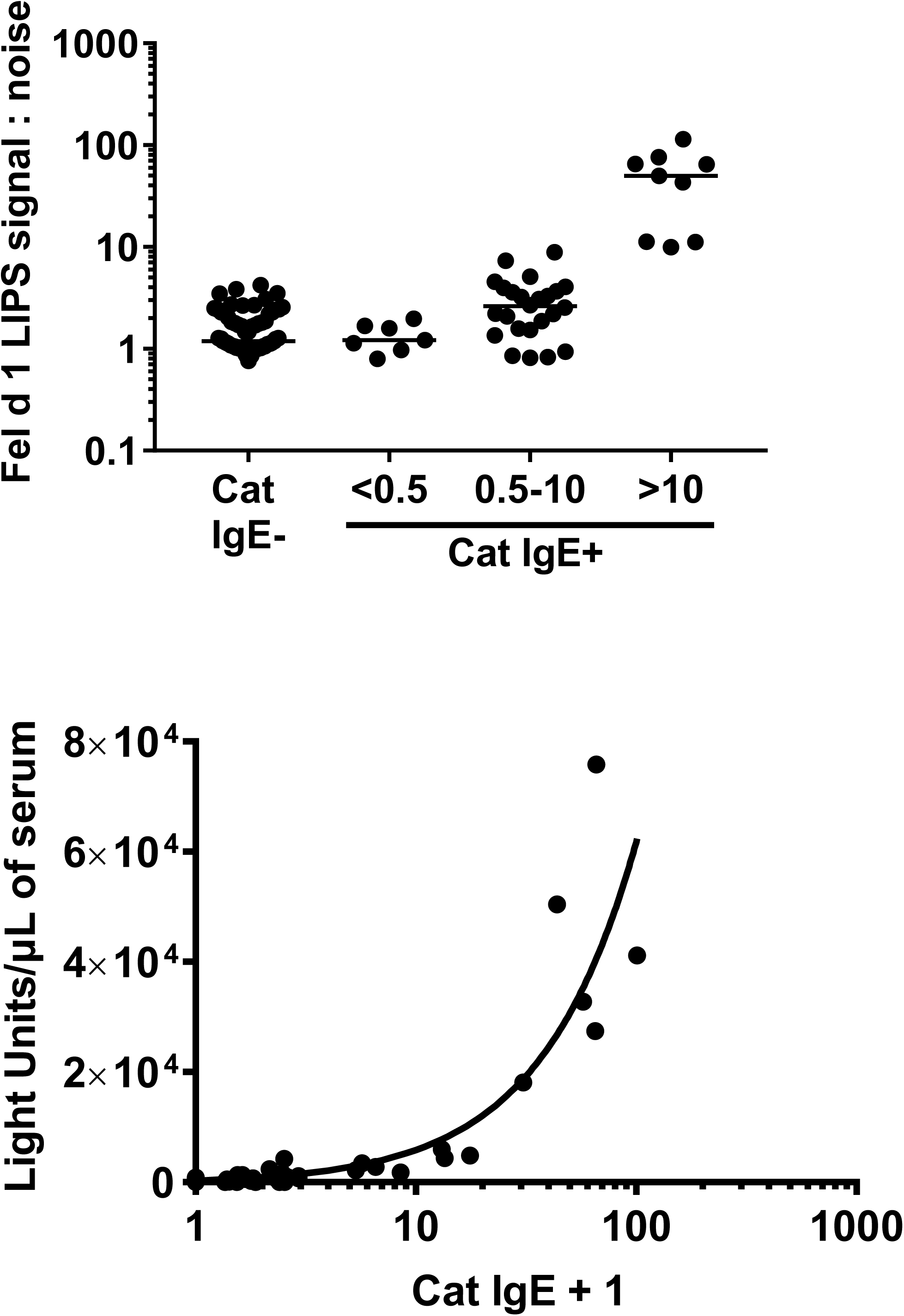
LIPS assay detects Fel d 1-specific IgE. Top panel: Fel d 1 LIPS signal: noise ratio in cat IgE- (*n* = 74) and cat IgE+ individuals, subdivided into cat IgE < 0.5 kU/mL (*n* = 7), 0.5-10 kU/mL (*n* = 24), and > 10 kU/mL (*n* = 9). Significant differences were found amongst all groups (Kruskal-Wallis test) and between all pairs except for cat IgE- and cat IgE+ < 0.5 (Mann-Whitney test). Bottom panel: Light units per µl of serum vs. cat IgE + 1 (to accommodate log scale). *n* = 91.

## Discussion

These results demonstrate that LIPS immunoassays can quantify component-specific IgE in small volumes of human serum. Although our sample cohort was relatively small and heterogeneous, the sensitivity and specificity of the LIPS assay for identifying cat IgE was similar to that reported for aeroallergen skin prick testing.^4^ We did not have information regarding skin prick sensitivity to cat or clinical symptoms following cat exposure for our cohort; it is possible that the efficacy of LIPS would be increased if our analysis had been able to include these parameters.

For Fel d 1, the LU correlated with cat IgE levels by ImmunoCAP. This is consistent with studies showing that Fel d 1 is a reliable marker for cat sensitization and has high sensitivity and specificity for clinical reactivity to cat.^5^ For patients with positive cat IgE but low LIPS signal, it is either because their IgE specific for Fel d 1 is below the limits of detection of the assay (possible, but unlikely), or rather their IgE response is specific for a different Fel d component. Most other aeroallergens do not have a component with the same dominance over others that Fel d 1 has in cat;^2^ thus, using a single component to determine sensitivity may not be clinically useful. However, LIPS is a high throughput system where responses to multiple target antigens can be performed,^6^ and components could be combined in a single assay to more accurately determine allergen sensitivity. Indeed, we have piloted detection of IgE specific for 5 peanut components (data not shown).

The LIPS immunoassay offers a sensitive and efficient method to both qualitatively and quantitatively determine an individual’s sensitization patterns to major components of allergens, using substantially less serum than is typically sent for a single allergen-specific IgE. This is of importance in vulnerable populations, such as children or individuals with anemia or reduced cardiac output, who require evaluation for multiple allergens and allergen components and have limitations on blood collection volumes. LIPS could have major utility in food allergy, where component-resolved diagnostics are clinically available and are applied in determining likelihood of clinical reactivity.^7^ With increasing interest in the development of component-resolved immunotherapy, one can also envision LIPS as part of the process in creating customizable immunotherapy based on an individual’s sensitization profile.

In addition to detection of specific IgE, LIPS may also be used to determine levels of other antibody isotypes, such as IgG4 or IgA, specific to a particular allergen or allergen component, simply by changing the isotype specificity of the antibody used to precipitate antibody-protein complexes.^8^ The LIPS assay may also be modified to evaluate for quantification of components themselves,^9^ which would have utility in determining component levels in foods, environmental samples, and allergen extracts.

In summary, due to their sensitivity, use of small amounts of sample, and rapidity, LIPS immunoassays can be a powerful and malleable tool with numerous uses in allergy research, diagnostics, and treatment.

## Abbreviations/acronyms

LIPS: Luciferase immunoprecipitation systems;
Ruc: *Renilla* luciferase;
LU: Light units;
ROC: Receiver operating characteristic

